# Learning-related population dynamics in the auditory thalamus

**DOI:** 10.1101/2020.03.31.017632

**Authors:** Ariel Gilad, Ido Maor, Adi Mizrahi

**Affiliations:** Department of Medical Neurobiology, Institute for Medical Research Israel Canada, Faculty of Medicine, The Hebrew University, Jerusalem, 9112001, Israel; Department of Neurobiology, The Hebrew University of Jerusalem, Jerusalem, 91904, Israel; The Edmond and Lily Safra Center for Brain Sciences, The Hebrew University of Jerusalem, Jerusalem, 91904, Israel

## Abstract

Learning to associate sensory stimuli with a chosen action has been classically attributed to the cortex. Whether the thalamus, considered mainly as an upstream area relative to cortex, encodes learning-related information is still largely unknown. We studied learning-related activity in the dorsal and medial regions of the medial geniculate body (MGB), part of the non-lemniscal auditory pathway. Using fiber photometry, we continuously imaged population calcium dynamics as mice learned a go/no-go auditory discrimination task. The MGB was tuned to frequency shortly after stimulus onset and responded to cognitive features like the choice of the mouse several hundred milliseconds later. Encoding of choice in the MGB increased with learning, and was highly correlated with the learning curves of the mice. MGB also encoded motor parameters of the mouse during the task. These results provide evidence that the MGB encodes task- motor- and learning-related information.

## Introduction

The thalamus was long considered a passive relay station of sensory information to cortex. However, recent evidence suggests that thalamic nuclei may also be involved in high-order processing of cognitive functions such as attention, working memory and learning (Acsády, 2017; Audette et al., 2019; Bennett et al., 2019; Bolkan et al., 2017; Guo et al., 2017; McAlonan et al., 2008; Rose and Bonhoeffer, 2018; Roth et al., 2016; Saalmann and Kastner, 2015; Schmitt et al., 2017; Ward, 2013; Williams and Holtmaat, 2019; Zhang and Bruno, 2019). Despite this large body of evidence and partly due to the dozens of different thalamic nuclei (Jones, 1985), the role of specific thalamic nuclei in cognitive processing remains unclear. In this study we focused on the neural correlates of higher-order thalamic nuclei of the medial geniculate body (MGB; auditory thalamus) during auditory learning.

The MGB is the thalamic relay center of the auditory pathway, predominantly receiving direct input from the inferior colliculus (Calford and Aitkin, 1983; Peruzzi et al., 1997), but also from cortex (Winer et al., 2001) and other sources (Crabtree, 1998; Lee, 2015; Winer, 1992). Its projections target the cerebral cortex and numerous other brain areas such as the amygdala (LeDoux et al., 1991; Lee, 2015). Anatomical studies divide the MGB into three main sub-divisions: ventral, dorsal and medial (Calford and Aitkin, 1983; Clerici and Coleman, 1990; Hashikawa et al., 1991; Imig and Morel, 1985; Mo and Sherman, 2019; Rouiller et al., 1989; Smith et al., 2012). The ventral MGB relays topographically organized information from the inferior colliculus to the primary auditory cortex (Rouiller et al., 1989; Smith et al., 2012). This pathway is called the lemniscal pathway, and is considered to be the main auditory processing pathway. In contrast, the dorsal and medial parts of the MGB project to higher-order cortical areas (Huang and Winer, 2000a; Lee, 2015) and receive cortical feedback, among others, from layer 5 of the auditory cortex (Bartlett et al., 2000a; Lee, 2015; Llano and Sherman, 2008a). Thus, the medial and dorsal parts of the MGB, considered to be part of the non-lemniscal pathway, are well positioned to encode higher-order information of sensory, motor and associative nature.

Learning, the process of acquiring new knowledge through experience, is traditionally thought to involve the neocortex. Learning to discriminate between different stimuli leads to changes in the respective primary sensory areas (Blake et al., 2002; Chen et al., 2015; Driscoll et al., 2017; Gilad and Helmchen, 2019; Jurjut et al., 2017; Komiyama et al., 2010; Li et al., 2008; Makino and Komiyama, 2015; Poort et al., 2015; Yan et al., 2014). Neural responses have often been shown to strengthen after learning. For example, neural signals in animals that gain expertise in a specific task, show increased activity and result in higher discriminatory power between the learned stimuli (Gilad et al., 2018; Gilad and Helmchen, 2019; Poort et al., 2015; Wiest et al., 2010; Yan et al., 2014). In the auditory system, when training to discriminate between two sounds, neurons in auditory cortex display several learning related modulations: encoding of higher-order choice information (Guo et al., 2019; Jaramillo et al., 2014), tonotopic map reorganization (Maor et al., 2020; Polley et al., 2006), increased responses to a target frequency and decreased responses to a distractor stimulus (Blake et al., 2002; David et al., 2012; Ghose, 2004; Ohl and Scheich, 2005). In addition, a large body of evidence indicates that the medial MGB is involved in auditory fear conditioning, by providing fast, less refined auditory information to the lateral amygdala (Han et al., 2008; Herry and Johansen, 2014; Maren and Quirk, 2004; Quirk et al., 1995; Romanski and LeDoux, 1992; Weinberger, 2011), an indication of experience dependent synaptic plasticity in the MGB. Nevertheless, whether MGB encodes learning-related modulation beyond the fear system, remains unknown (Chen et al., 2019; Jaramillo et al., 2014). To study learning related modulations in the MGB, we chronically imaged MGB population responses to sounds as mice learned a go/no-go auditory discrimination task. We find learning-related modulations in the MGB, and a particularly strong modulation to choice signals.

## Results

### Calcium imaging from the MGB along learning

To study learning-related changes in the MGB we first injected AAV-GCaMP6f into the MGB of C57BL/6 mice and implanted a 400 µm optical fiber directly above the injection site. After a week of handling and habituation to head-fixation, we trained mice on a go/no-go auditory discrimination task. Each trial started with a visual start cue (orange LED; duration 0.1 s; 2 seconds before stimulus onset) followed by an auditory stimulus, either a go or a no-go pure tone sound (Fig. 1A; duration 1 second; sounds separated by 0.5 octave; mostly 10 kHz for go and 7.1 kHz for no-go; Methods). After stimulus offset, we counted licks in a virtual response window of 3 seconds. Licking in response to the go sound were counted as ‘hit’ trials and rewarded with a drop of water. Withholding licking in response to the no-go sound were counted as correct rejection trials (CR), and were not punished or reinforced. Licking in response to the no-go sound were counted as false alarm trials (FA) that were followed by a mild punishment of white noise (duration 3 seconds). Withholding licking for the go sound were counted as Miss trials, and were not punished. As mice learned to discriminate between the two sounds we continuously imaged population responses in the MGB in addition to monitoring their body movements during the task (Fig. 1B). We first imaged six mice across learning. The fiber tips of the six mice were reconstructed to the higher-order regions of the MGB (Fig. 1C; MGBd and MGBm, grouped as MGB for simplicity). Mice learned the task within 1200-2000 trials as assessed by d’ (defined as d’ = Z(Hit/(Hit+Miss)) – Z(FA/(FA+CR) where Z denotes the inverse of the cumulative distribution function; learning threshold was defined as d’=1; Fig. 1D, E). The learning curves with respect to the go and no-go trials shows that mice varied in performance and strategy (Fig S1). Learning the task took different forms: some mice increased their CR rate, others had a steep increase in hit rate, whereas others gradually increased both hit and CR rates. Thus, mice learned the task using a range of behavioral strategies.

**Figure 1.**
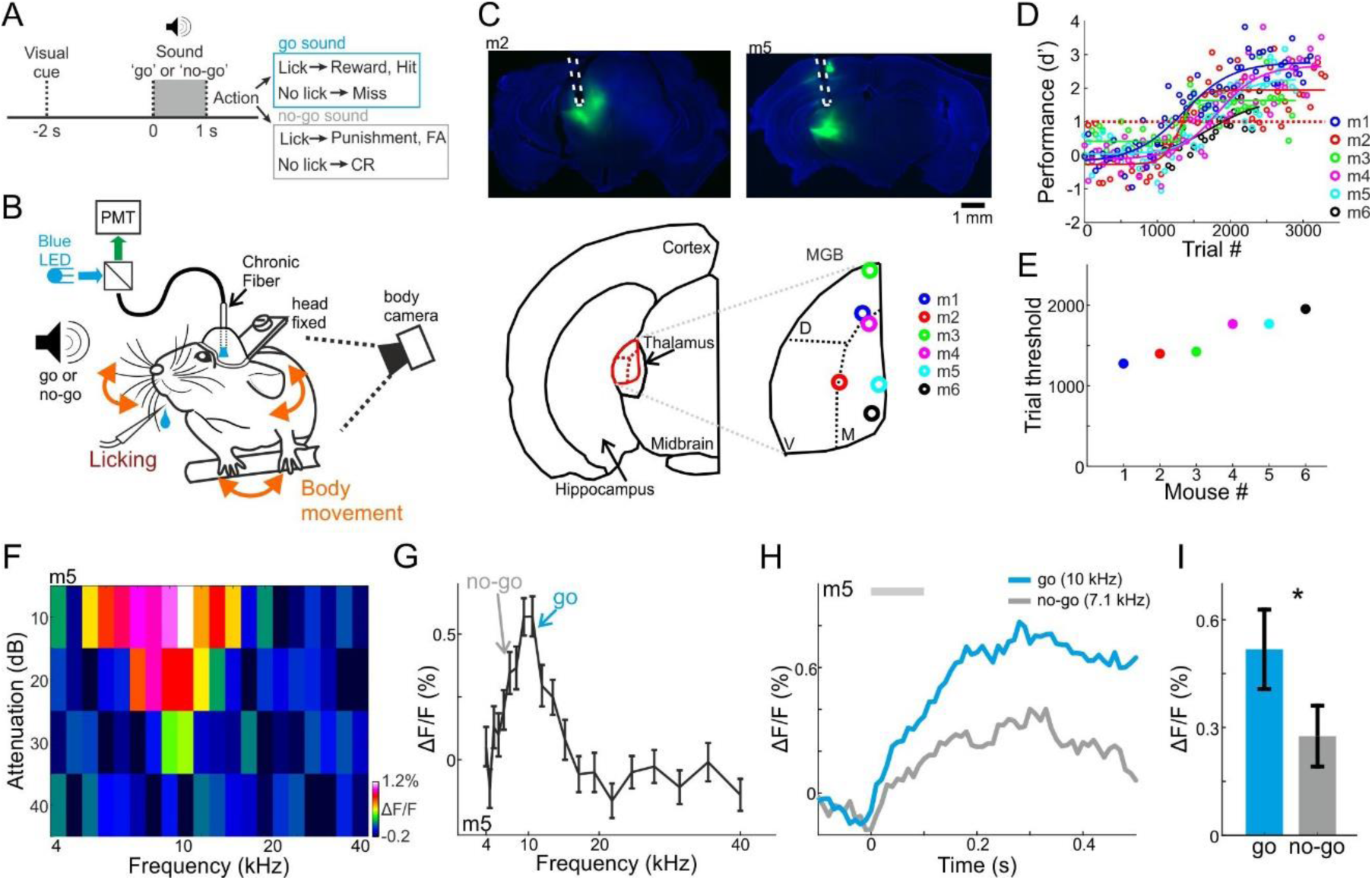
Behavioral paradigm, performance, and frequency tuning. **A.** Trial structure of a go/no-go auditory discrimination task and possible trial outcomes. **B.** Behavioral setup for head fixed behaving mice along with simultaneous fiber photometry in the medial geniculate body (MGB). **C.** *Top:* Fluorescent images of two coronal slices from two different mice, showing GCaMP6f (green) in the MGB along with the fiber track highlighted in white. Bottom: Localization of the fiber tip in the MGB for all 6 mice. **D.** Behavioral learning curves for all mice (n=6) depicting the performance (d’) as a function of trial number. Each learning curve was fitted with a sigmoid function. Dashed red line indicates the performance threshold (d’=1). **E.** Learning threshold (the trial number where the learning curve crossed the performance threshold) for all mice. **F.** An FRA plot (attenuation versus frequency) from one example recording of mouse #5. **G.** Frequency tuning curve for the same example mouse. The frequencies used as go and no-go are marked by arrows. **H.** Average responses to the go (blue) and no-go (gray) sounds. **I.** Average evoked response (calculated from the 100ms after stimulus onset; gray bar in ‘H’) to the go and no-go sounds from all mice. Error bars are s.e.m across mice. *P < 0.05. Wilcoxon sign rank test.

Prior to training each mouse was imaged while playing a range of frequencies (4-40 kHz; 10, 20, 30, 40 dB attenuations relative to 62 dB; duration 0.1 second; Methods). Population responses in the MGB displayed a classical V-shaped frequency response area (FRA, Fig. 1F). In the six mice shown below, the response to the go frequency was significantly higher as compared to the no-go frequency (Fig. 1G-H, one example mouse; Fig. 1I, average of all six mice; p<0.05; Signed rank test; Fig. S2A for the tuning curves of all mice). Since tuning curves of individual neurons in higher-order MGB (i.e. dorsal and medial parts) are generally broader than ventral MGB, the rather sharp tuning is surprising (Bordi and LeDoux, 1994; Weinberger, 2011). Fiber photometry is thought to record a relatively large population of neurons, and may be biased to the more sharply tuned population in MGB.

### MGB encodes the choice of the mouse

To evaluate how sounds and other task attributes were represented in the MGB across learning, we plotted responses to the go and no-go sounds as mice learned the task. Figure 2A shows responses to the go and no-go sounds of one representative mouse (Fig. 2A). The most evident change in MGB responses across learning was during the late part of the trial (0.6-1 seconds after stimulus onset), and particularly so for go trials (Fig. 2A).

**Figure 2.**
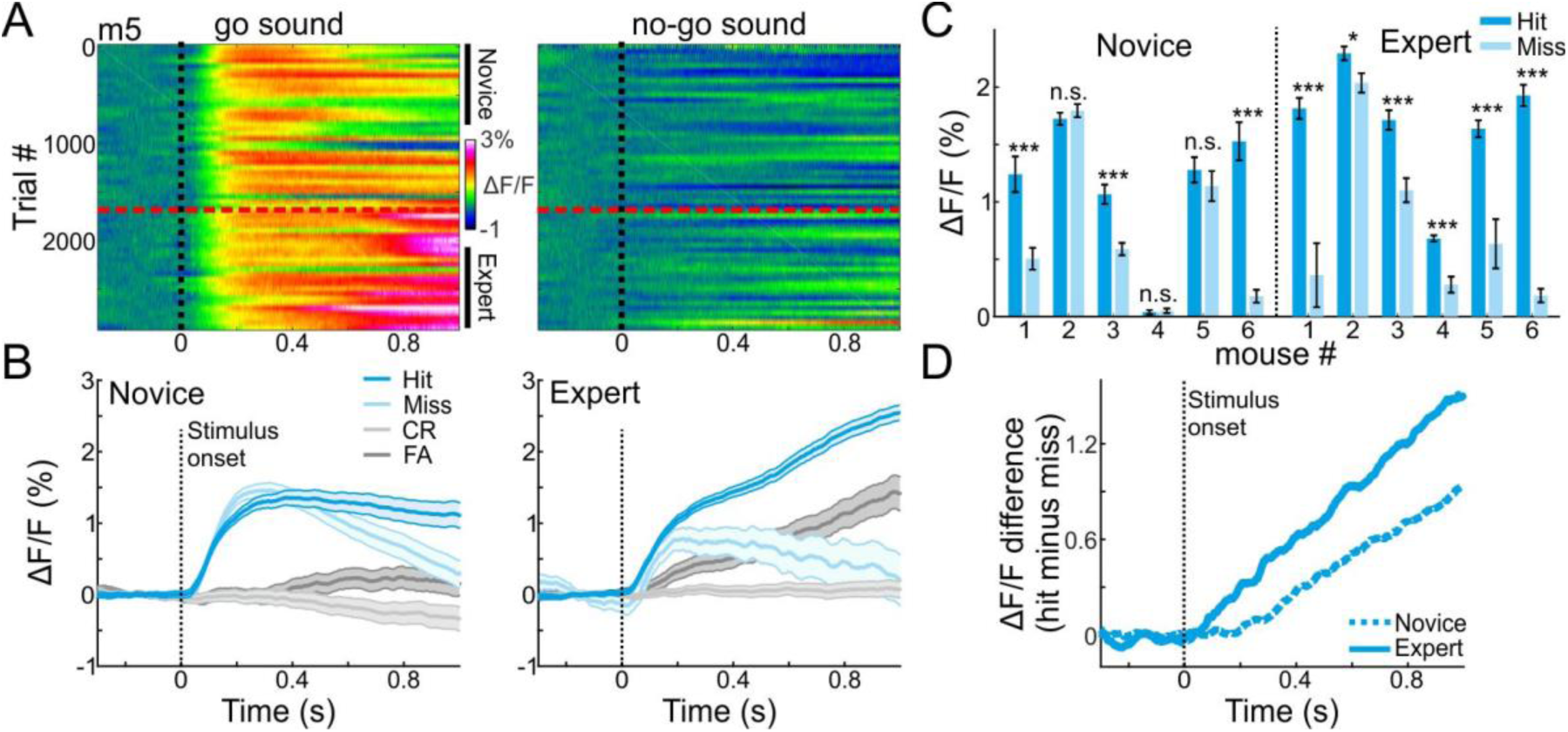
MGB encodes choice in expert mice. **A.** 2-dimensional plots of the calcium responses in MGB during the trial (x-axis) and across learning (y-axis; 50 trial bins) for one example mouse divided into go (left) and no-go (right) sounds. Dashed black line indicates stimulus onset (1 s duration) and dashed red line indicates the learning threshold. **B.** Calcium responses when the mouse was novice (left) and expert (right). Traces are shown separately for different trial types (hit, miss, CR and FA trials; same data as in ‘A’). Shaded error bars are s.e.m across trials. **C.** Mean calcium response during stimulus presentation per mouse in hit and miss trials when mice were novice (left) and expert (right). Error bars are s.e.m across trials. **D.** Choice responses (defined as the response difference between hit and miss trials) during the trial, averaged across all mice when they were novice (dashed line) and expert (solid line). *P < 0.05. ***P < 0.001. n.s. – not significant. Wilcoxon rank sum test.

A particularly informative comparison was between hit and miss trials, where the stimulus is identical but the choice of the mouse is different (either lick or no-lick). Thus, differences between hits and misses represent encoding of choice. In expert mice (defined as the last 500 trials), MGB responses were higher for hit as compared to miss trials, and more so than in the novice mice (defined as the first 500 trials; Fig. 2B; compare blue to light blue traces; For comparison, CR and FA trials are plotted in gray). Higher responses in hit versus miss trials were evident in all (6/6) expert mice and in 50% (3/6) of novice mice (Fig. 2C; p<0.05; Wilcoxon rank sum test for each mouse separately). Choice responses (i.e. hit minus miss) increased gradually after stimulus onset, and were stronger in expert mice (Fig. 2D). These data provide evidence that the MGB encodes more than only sounds. MGB encodes the choice of the mouse, which implies that the auditory thalamus is involved in higher-level sensory-motor processing or cognitive attributes of the task.

### MGB encodes sounds early and choices late

To further explore the role of MGB in high level processing versus low-level sound processing we tested the discrimination between sensory stimuli and choices at the single trial level. To do so, we calculated the receiver operating curve (ROC) and derived the area under the curve (AUC) between pairs of different trial-type distributions. We define three different AUC measures by comparing hit trials to either CR (task AUC; brown; i.e. different stimuli and different choices), to FA (stim AUC; magenta; i.e. different stimuli but similar choice) or to Miss (choice AUC; green; i.e. similar stimuli but different choice) trials (Fig. 3A). The AUC value ranges from 0 to 1 and quantifies the accuracy of an ideal observer. AUC values close to 0.5 indicate low discrimination whereas values away from 0.5 indicate high discrimination. Task-AUC is useful to describe the maximal discrimination of all task attributes, but cannot dissociate between the different stimuli or choices. Thus, the Stim- and Choice-AUCs complement the task-AUC. Together, these comparisons encompass the full breadth of options in the task (see below for additional trial types). In addition, we calculated AUC values for each time frame along the trial for the novice and expert conditions separately (i.e. first and last 500 trials). As a control, we calculated the sample distribution of trial-shuffled data and calculated the AUC identically. We define significance as the time point when the observed AUC values exceed ±3std above the mean of the shuffled distribution (dashed lines in Fig. 3B). For example, data from one mouse is shown in Figure 3B, where both ‘task-AUC’ and ‘stim-AUC’ values are significantly discriminative early in the trial in both the novice and expert cases (Fig. 3B, <100 ms from stimulus onset; arrows). This example shows that MGB responses carry highly discriminative information between the stimuli early in the trial and that this does not change following learning. In contrast, ‘choice-AUC’ reached significance only in the expert condition and only later in the trial (Fig. 3B, green trace, >200ms from stimulus onset; arrow).

**Figure 3.**
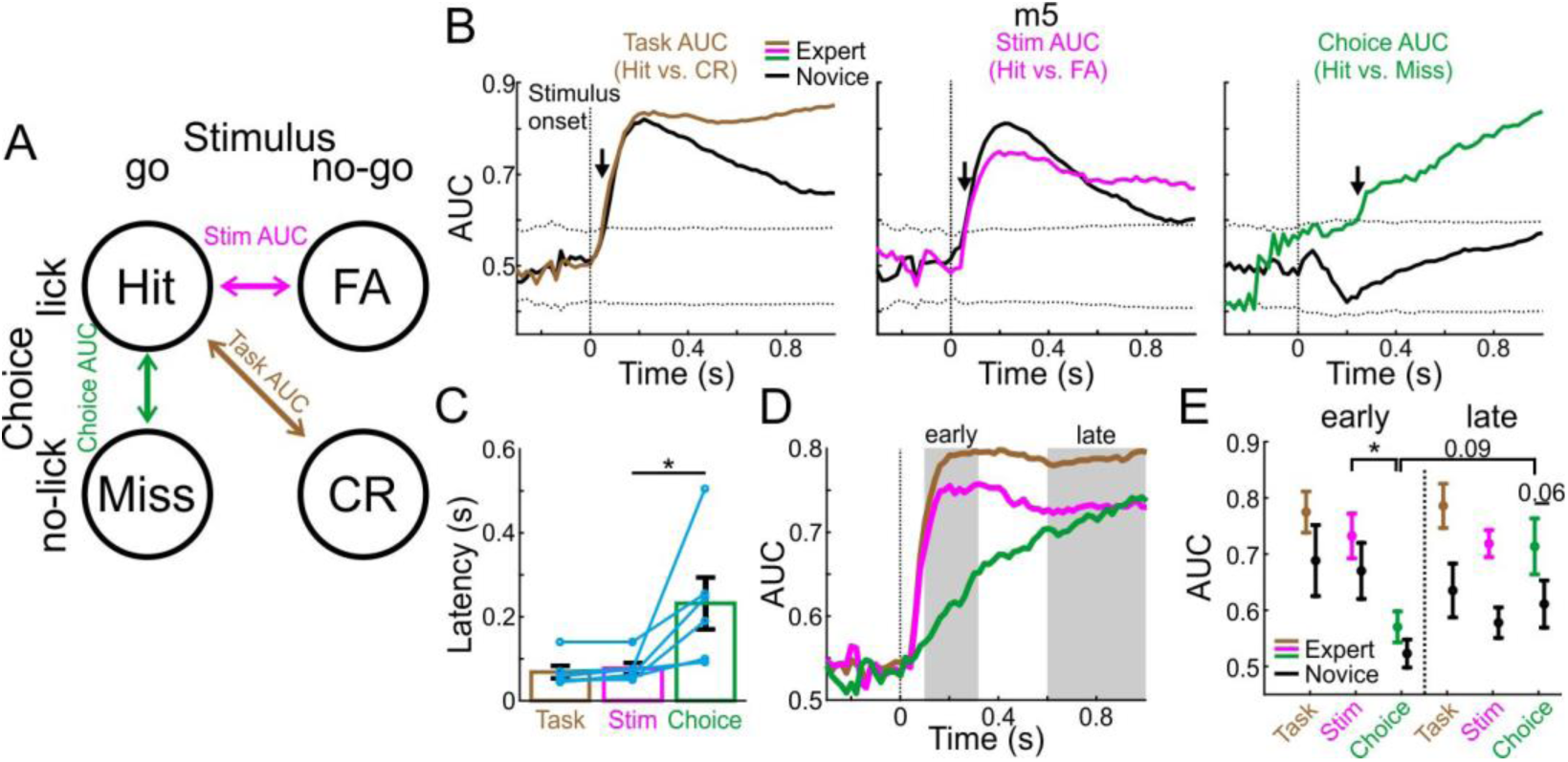
MGB discriminates early between stimuli and late between choices. **A.** Schematic of the three single trial discrimination measures: hit vs. CR (brown; Task AUC), hit vs. FA (magenta; Stim AUC) or hit vs. miss (green; Choice AUC). **B.** Task, Stim and Choice AUCs during the trial for one example mouse when it was novice (black lines) and expert (colored lines). Dashed black lines display the mean±3s.t.d of trials from shuffled data. Arrows indicate the latency of the AUC measure (i.e. the first time point the signal exceeded the shuffled data). **C.** Latency of discrimination for the Task, Stim and Choice AUCs for all mice (averaged and marked individually). Error bars are s.e.m across mice. **D.** Task, Stim and Choice AUCs averaged across all expert mice. Early and late times are marked in gray. **E.** The three AUC measures averaged during early (left) and late (right) times when mice were novice (black lines) and expert (colored). Error bars are s.e.m across mice. *P < 0.05. Wilcoxon signed-rank test.

To quantify this effect across mice, we defined the latency for discrimination as the time it takes to reach significance for each AUC measure in the expert case (arrows in Fig. 3B). The latency of discrimination for ‘stim-AUC’ was significantly lower than that of the ‘choice-AUC’ (Fig. 3C; p<0.05; Signed rank test). When we averaged across all mice, task- and stim-AUCs increased early and abruptly whereas choice-AUC increased gradually (Fig. 3D; choice-AUCs for individual mice are shown separately in Fig. S3). We then pooled together the AUC values during the early (0.1-0.3 seconds after stimulus onset) and late (0.6-1 seconds after stimulus onset) times of the trial (Fig. 3E). We found that 1) Stim-AUC was significantly higher than choice-AUC early in the trial (p<0.05; Signed rank test), 2) Choice-AUC was higher, yet not significantly, during the late part of the trial compared to early times (p=0.09; Signed rank test), and 3) Choice-AUC was higher, yet not significantly, in the expert compared to the novice state, particularly during late times during the trial (Fig. 3E; p=0.06; Signed rank test). In addition, we calculated choice and stim AUCs based on other pairs of trial types: CR vs. Miss for an alternative stim-AUC and FA vs. CR for an alternative choice-AUC (Fig. S4). Stim-AUC displayed similar early onset discrimination in both naïve and expert cases. In contrast, choice-AUC did not show significant discrimination in some mice. This was mainly due to the fact that the responses were quite weak to the no-go sound, resulting in a low discrimination value. In summary, these data indicate that mostly choice encoding changes with learning and that this information develops late - several hundred milliseconds after stimulus onset.

### Plasticity in MGB correlates with learning

Continuous imaging throughout the whole learning process enabled us to investigate the activity of the MGB on a trial-by-trial manner, and compare it with behavioral performance of each mouse. For this analysis, which was focused on go trials only, we averaged the calcium signals at late times during the trial (0.6-1 seconds after stimulus onset) to obtain MGB responses as a function of learning (termed ‘MGB response curve’). The MGB response curve was sigmoid-like, resembling the learning curve of the mouse (Fig. 4A shows data from two mice; line is a sigmoid function fit). Next, we calculated the MGB response curve for each time point along the trial separately, correlated it with the learning curve, and plotted the correlation coefficient along time (see Methods). Positive correlations depict high similarity between the MGB response curve to the learning curve at a given time point. The correlation increased after stimulus onset, indicating a strong relationship between MGB modulations and the learning process (Fig. 4B - two representative mice; Fig. 4C-all mice). To find the maximum point of change for each MGB response curve, we normalized sigmoid fits of the MGB response curves for each mouse separately, and derived the trial threshold for each curve (Fig. 4D; defined as the trial number in which the normalized curve crosses 0.5). The trial threshold of the MGB curves matched nicely the learning threshold of each mouse (Fig. 4E; r=0.89; p<0.05). To test whether MGB responses had undergone changes not directly related to learning, we compared between the responses to tones in passively listening mice before and after mice learned the task. We found no differences between the MGB frequency tuning curves or the response to the go frequency during passive listening, before and after learning (Fig. S5), indicating that MGB plasticity occurs specifically for task related signals, at least at the population level. In summary, we found that changes in MGB responses are positively correlated with learning as evident in similar modulation of the choice signal and the performance of each mouse separately.

**Figure 4.**
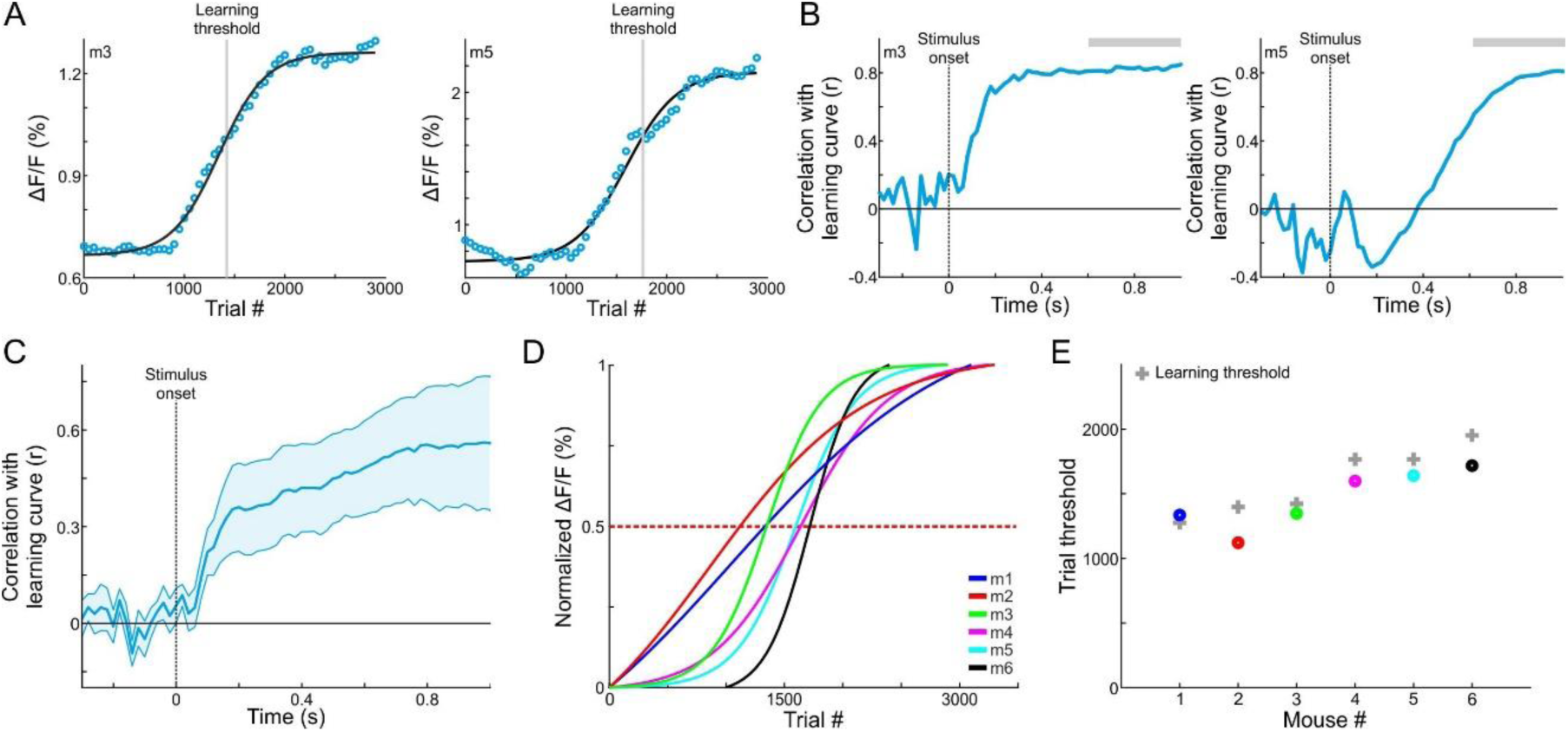
MGB changes correspond with gaining expertise. **A.** MGB responses (averaged during late times, 0.6-1 seconds after stimulus onset; gray bars in B) as a function of learning in two example mice. Each MGB curve was fitted with a sigmoid function (black lines). The learning thresholds are marked with a gray line. **B.** Correlation coefficient as a function of time between the MGB curve and the learning curve of the two example mice shown in ‘A’. Positive values indicate that MGB responses change in a similar way to the behavioral performance of the mouse. **C.** MGB responses correlate with learning. The trace is the average correlation across all 6 mice. Error bars are s.e.m across mice. **D.** Normalized MGB curve fits of all mice. Dashed red line indicates threshold. **E.** Maximal change in MBG responses (i.e. the trial number in which the MGB fit crossed 0.5, dots) as a function of learning threshold (gray crosses).

### Body movements affect MGB responses

The strong relationship between MGB plasticity and learning prompted us to further investigate other parameters that may affect learning-related modulations, specifically related to encoding of choice. One factor that may affect MGB responses are parameters related to movement (e.g. forelimb movement, nose twitching, whisking, licking etc.), which were recently found to have substantial impact on neuronal activity in different brain areas (Gilad et al., 2018; Gilad and Helmchen, 2019; Musall et al., 2019; Stringer et al., 2019). As mentioned above, we continuously monitored the body movements of the mouse throughout the experiment (Fig. 1A, 5A). As expected, changes in body movements were correlated with learning. As soon as mice crossed the learning threshold, they exhibit more body movements as evident by plotting their movement probability (Fig. 5A, right). Movement probability was high both within the stimulus period (0-1 seconds after stimulus onset) and, as expected, during the response period (Fig. 5A, B). Movement probability during stimulus presentation across learning was sigmoid-like, and significantly higher in experts compared to novice mice (Fig. 5C, D; p<0.05; Signed rank test). Thus, as part of the development of choice responses in MGB, the body movements of mice during the task were also strongly related to the learning process.

**Figure 5.**
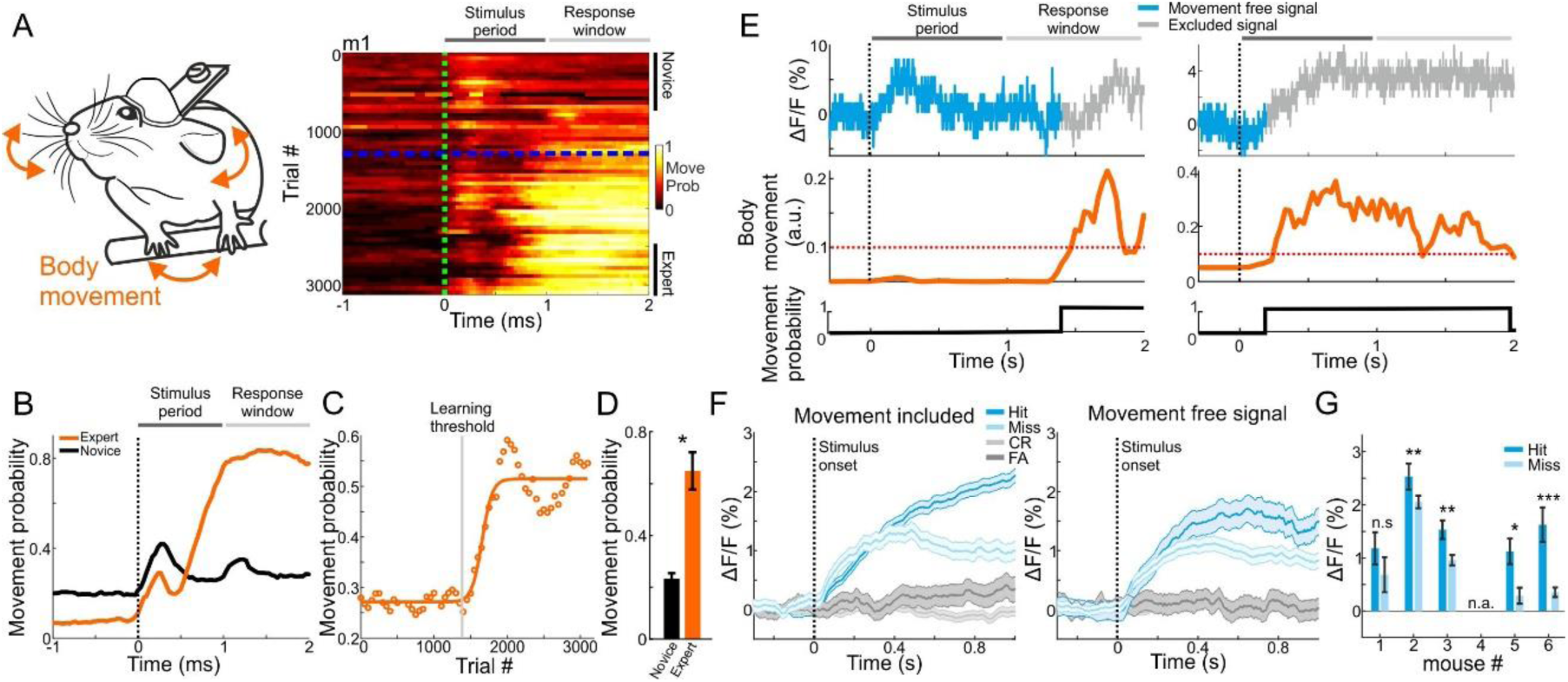
Body movements affect MGB responses. **A.** Left: schematic illustration of mouse body movements during the task. Right: A 2-dimensional plot of movement probability during go trials within the trial (x-axis) and across learning (y-axis; 50 trial bins) for an example mouse. Dashed green line indicates stimulus onset (1 s duration) and dashed blue line indicates learning threshold. Color scale bar indicates min/max of movement probability. **B.** Movement probability during the trial for the example mouse in A when it was novice (black) and expert (orange). **C.** Movement probability (averaged during late times; 0.6-1 sec after stimulus onset; go trials) across learning fitted with a sigmoid function. Same data as in ‘B’. **D.** Movement probability during late times for novice and expert averaged across all mice. Error bars are s.e.m across mice. *P < 0.05. Wilcoxon signed-rank test. **E.** Single trial traces of MGB calcium response (top), body movement (middle) and a binary movement vector (bottom; thresholding the body movement, dashed red line; Methods). To exclude effects of body movements, MGB responses were truncated on a trial-by-trial basis. **F.** MGB calcium responses for different trial types when including (left) and excluding (right) movement. Data is from one example mouse (same as Fig. 3B). Error bars are s.e.m. **G.** Mean MGB responses (averaged during stimulus presentation) for movement-free hit and miss trials per mouse. F, G are data from expert mice. Error bars are s.e.m across trials. *P < 0.05. **P < 0.01. ***P < 0.001. n.s. – not significant. Wilcoxon rank sum test.

The strong effects of movement suggest that the abovementioned MGB ‘choice’ signal may simply represent body movements rather than choice, *per se*. Indeed, some MGB responses and body movement covaried strongly whereas others did not (Fig. 5E). Thus, to rule out the possibility that movement is the dominant feature in our calcium signal, we calculated ‘movement free’ MGB responses by detecting the movement onset in each trial and truncating the MGB signal when mice moved (Methods; Gray traces in Figure 5E). This allowed us to evaluate separately MGB responses without the body movements. Movement-free MGB responses still encoded the choice of the mouse. Specifically, we found higher responses in hit compared to miss trials (Fig. 5F). Notably, the effect size was smaller and levels of statistical significance were weaker (Fig. 5G; compare to Fig. 2C ‘Expert’). Importantly, when mice were still novice, movement-free MGB responses did not encode choice, indicating that choice encoding develops with learning (Fig. S6). To rule out movement-related artifacts, we also imaged mice during the task while exciting the MGB with a wavelength that does not excite the calcium indicator (565 nm). We found no movement related artifacts (data not shown). In summary, body movements during the task are evident in the calcium signal and are learning-related. Nevertheless, MGB responses still maintains choice information that is separate from the motor parameters and, critically, develops with learning. These data strengthen the claim the MGB encodes high-level information.

### Learning related changes in MGB differ in different frequency bands

As noted above, MGB responses were tuned to frequency (Fig. 1F-I). We next tested whether the changes we observed in MGB are general or specific to the response properties at the frequencies we used. To this end, we trained three additional mice on the task. In these mice, the go frequency did not correspond to the best frequency, i.e. the underlying MGB population response was not highest for the target frequency (tuned away from the go frequency; Fig. 6A shows one mouse with the go signal in the outskirts of the tuning curve; Fig. S2B for the tuning curves of all three mice). Interestingly, MGB responses in all three mice still showed choice encoding but the absolute direction of change was in the opposite direction as compared to the first cohort of mice (compare Fig 6B to 2C). Specifically, the MGB responses to the hit trials was now significantly lower than to the miss trials and this was evident in expert mice only (Fig. 6B; p<0.001; Signed rank test for each mouse separately). In expert mice, choice encoding gradually developed with time along the trial and was evident only in expert mice (Fig. 6C). Here again, MGB response curves were sigmoid-like (Fig. 6D) and strongly correlated with learning (Fig. 6E). A more detailed description of these effects is shown in Figure S7. Taken together, we infer that learning enhances responses to the target frequency while simultaneously suppressing activity away from the target frequency. Notably, both phenomena still result in the same effect (i.e. encoding of choice), but exploit different mechanisms, as we discuss below.

**Figure 6.**
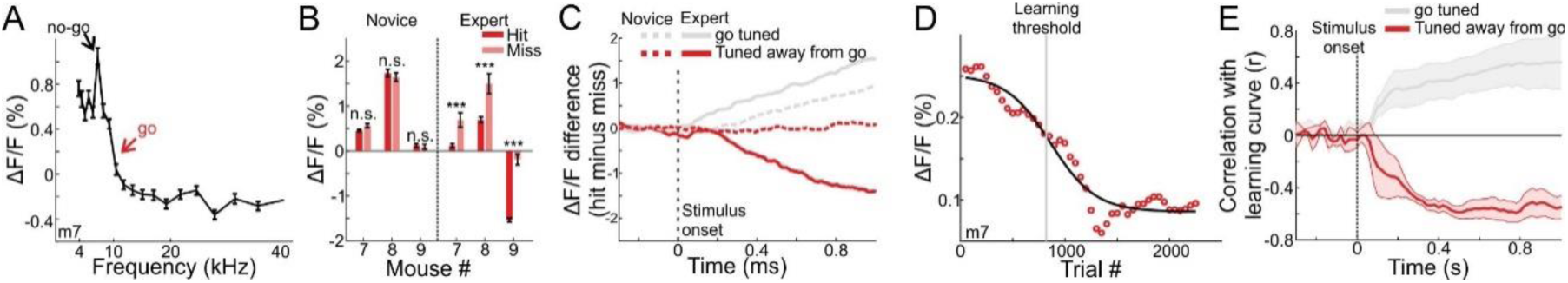
MGB responses tuned away from the go frequency are suppressed during learning. **A.** Frequency tuning curve for an example mouse with peak tuning frequency away from the go sound (tuned away from go). **B.** Mean calcium response during stimulus presentation per mouse in hit and miss trials. Left-novice mice, Right-expert mice. Error bars are s.e.m across trials. ***P < 0.001. n.s. – not significant. Wilcoxon rank sum test. **C.** Choice responses, averaged across all three mice that are tuned away from the go frequency, for the expert (solid line) and novice (dashed line) conditions. Gray traces are for the go-tuned mice, same as Fig. 2D. **D.** MGB response curve along learning in one mouse that is tuned away from the go frequency (compare to Fig. 4A). **E.** Correlation between MGB response curves and learning curves as a function of time, averaged across the 3 mice. Error bars are s.e.m across mice. Gray traces are for the go-tuned mice, same as Fig. 4C.

## Discussion

We describe learning-related changes in evoked responses to sounds in the auditory thalamus. We find that auditory thalamus encodes the choice of the mouse several hundred milliseconds (~200 ms) after sound onset, by specifically enhancing activity to the go frequency as learning proceeds. Thus, late thalamic responses display different dynamics for the same sound depending on its learned state. The medial and dorsal parts of the MGB are part of a thalamocortical loop that has been suggested to encode high-order cognitive information (Bolkan et al., 2017; Guo et al., 2017; Schmitt et al., 2017). Choice encoding in the thalamus (Chen et al., 2019; Gimenez et al., 2015; Jaramillo et al., 2014) bears similarity to previous observations in the cortex (Gilad et al., 2018; Guo et al., 2019; Harvey et al., 2012; Pho et al., 2018; Siegel et al., 2015; Yang et al., 2015), indicating the involvement of the thalamocortical loop during learning. Choice encoding was consistent across mice and evident at the population level. Since we are unable to dissect the fiber photometry signal to its single cell components, we can only speculate about the nature of cells encoding choice. The increase in choice signal could arise from a large fraction of MGB neurons encoding choice after learning or from a smaller population that strengthens its responses. A previous study found that neurons in the MGB mediate choice to a limited extent, whereas only a small fraction (16%) of recorded neurons encoded choice information, supporting the latter possibility (Jaramillo et al., 2014). Notably, however, Jaramillo et al. (2014) recorded responses from all MGB sub-nuclei, including ventral MGB which might not encode choice, and thus underestimate the contribution of the non-lemniscal pathway to signal choice. In contrast, we targeted only the medial and dorsal MGB, thus, dissociating more efficiently between lemniscal and non-lemniscal thalamic nuclei (Chen et al., 2019). Moreover, further functional differences between these two subdivisions is expected. For example, we expect that the dorsal MGB will be affected more prominently by motor parameters compared to the medial MGB (Bartlett et al., 2000b; Huang and Winer, 2000b; Lee, 2015). As mentioned above, fiber photometry lacks single cell resolution and our conclusion holds at the population level only. It will be interesting to test which sub-population of MGB cells encode different task-related information. For example, specific populations of MGB neurons that project to different targets, likely transfer specific learning-related information in a selective manner (Chen et al., 2019).

Using chronic fiber photometry, we were able to continuously image MGB populations on a trial-by-trial manner as the mouse learned (Gilad and Helmchen, 2019). Interestingly, MGB displayed a sigmoid-like activity curve in response to the same sound, where the most abrupt MGB change coincided with the learning threshold of the mouse (i.e. when the mouse gained expertise). This simultaneous change in activity and learning is similar to the observed change in cortical areas (Gilad and Helmchen, 2019) and may indicate a tight relation between learning, thalamus and cortex. Our results imply that choice information may already originate in MGB, which is then relayed to the auditory cortex (Guo et al., 2019). Alternatively, the MGB may be activated via the thalamocortical loop with auditory cortex. Simultaneous recordings with tight regulation of the information transfer between thalamus and cortex will be instrumental in deciphering the origin of choice information. Nevertheless, MGB responses alone allowed us to predict with high precision when (across trials) did each mouse learn the task. Taken together, the trial-by-trial nature of our analyses provides strong evidence in relating between MGB plasticity and the learning process.

The observed plasticity in MGB points to pronounced plasticity during the training phase, that may involve several factors such as inhibitory effects, excitation-inhibition balance, synaptic plasticity or top-down interactions (Audette et al., 2019; McAlonan et al., 2008; Rose and Bonhoeffer, 2018). Learning-related dynamics may involve other circuit elements such as deep cortical layers (Audette et al., 2019; Olsen et al., 2012), inhibitory subtypes (Hashikawa et al., 1991; Khan et al., 2018; Peruzzi et al., 1997) or other thalamic sub-nuclei (Audette et al., 2019; Bennett et al., 2019). Indeed, higher-order thalamocortical connections were recently shown to drive synaptic plasticity in cortex during learning (Audette et al., 2019) and this may reciprocally trigger plasticity in the thalamus via corticothalamic feedback. The prolonged latency of choice development in our study, further implies an involvement of a late top-down interaction. An interesting sub-thalamic nucleus that may be involved in gating higher-order information is the thalamic reticular nucleus (TRN) which sends GABAergic projections to MGB (Crabtree, 1998; Lee, 2015). TRN could gate incoming sensory information, where in the novice case it’s expected to exert strong inhibition to MGB which is then decreased with learning; thus enabling enhanced activation to the go sound already at the level of the MGB (McAlonan et al., 2008; Nakajima et al., 2019). Such factors may also contribute to the enhanced discrimination between go and no-go trials in the MGB for expert mice, in which the above mechanisms may drive population enhancement in MGB specifically for ‘go’ sounds. In addition to enhancement in the go band of the MGB, we also observed a suppression in MGB to sounds that are tuned away from the go frequency. This phenomenon may be similar to visual surround suppression found in visual thalamus in which non-optimal features are suppressed, possibly by cortical feedback or lateral inhibition (Fisher et al., 2017; Jones et al., 2012; Wilke et al., 2009). A simultaneous enhancement/suppression scheme already in the MGB, may make the target frequency, (associated with a future reward) more salient (Nakajima et al., 2019).

We find that motor parameters, such as body movements, are also learning-related. As mice learn to associate between the go sound and a reward, they increase their body movement in preparation for licking and the upcoming reward, well within the stimulation period. MGB responses were also affected by body movements to different extents, as might be expected by the motor efferent cortical feedback from layer 5 (Bartlett et al., 2000b; Lee, 2015; Llano and Sherman, 2008b). Another possibility is that movement related encoding may ascend from the inferior colliculus, an upstream area (Gruters and Groh, 2012; Yang et al., 2020). The fact that MGB still encodes choice after truncating movement signals, indicate that the MGB does not inherit all task information from the inferior colliculus, but rather integrates information coming from both top-down and bottom-up projections. The effect of motor parameters on neuronal signals were recently observed in a brain-wide manner, emphasizing the need to strictly monitor and consider motor parameters during different tasks (Gilad et al., 2018; Gilad and Helmchen, 2019; Musall et al., 2019; Stringer et al., 2019); and particularly so when measuring high order information. In summary, auditory thalamus is shown to encode higher-order information during the time course of learning, implying that the learning process requires brain-wide activity spanning across both cortex and sub-cortex.

## Acknowledgements

We thank Joseph Jubran for help with the programming of the behavioral system. We thank members of the Mizrahi laboratory for commenting on early versions of this manuscript. This work was supported by an ERC consolidators grant to A.M. (#616063), Israeli Science Foundation grants to A.M. (#224/17, #2453/18), and the European Union’s Horizon 2020 research and innovation programme under the Marie Skłodowska-Curie grant agreement No 659719 to A.G.

## Author contributions

A.G., and A.M. designed the experiments. I.M. built the behavioral apparatus and integrated auditory systems. A.G. conducted the experiments and analyzed the data. A.G and A.M. wrote the paper.

## Data Availability

The data that support the findings of this study are publicly available at: https://osf.io/mt3bc/.

## Materials and Methods

### Animals

A total of n=9, 8-16 week-old female C57BL/6 mice were used in this work. All experiments were approved by Institutional Animal Care and Use Committee (IACUC) at the Hebrew University of Jerusalem, Israel.

### Surgery

To express a calcium indicator in MGB neurons and implant an optical fiber for imaging, mice were anesthetized with 2% isoflurane (in pure O2) and body temperature was maintained at 37°C. We applied local anesthesia to the area of surgery (lidocaine 1%), exposed and cleaned the skull. Next, we drilled a small hole in the skull and injected ??nl of an AAV virus pAAV.Syn.GCaMP6f.WPRE.SV40 (AAV9; from Addgene) using a glass pipette, to the dorsal or medial part of the MGB (3.3 mm posterior to bregma; 1.9 mm lateral to bregma; 3 mm in depth). Using the exact same coordinates, we then inserted a 400 µm optical fiber with an attached cannula (CFMC14L05; Thorlabs), directly above the injection site (2.95 mm deep). We chronically fixed the fiber position using dental cement that was mixed with a few drops of superglue for extra strength and stability. Finally, a metal post for head fixation was glued onto the bone in the back side of the right hemisphere. This procedure enabled a chronic and reliable imaging of population calcium responses from the MGB.

#### Auditory discrimination task

Mice were trained on a go/no-go auditory discrimination task (Fig. 1A). Each trial started with a visual start cue (orange LED placed in front of the mouse; duration 0.1 s; 2 seconds before stimulus onset) followed by an auditory stimulus, either a go or a no-go pure tone (Fig. 1A; duration 1 second; 62 dB; sounds separated by 0.5 octave). All mice were trained on 10 kHz for go and 7.1 kHz for no-go except for one mouse which was trained on 8.2 kHz for go and 5.8 kHz for the no-go. After stimulus offset, we counted licks in a virtual response window of 3 seconds. Licking in response to the go sound were counted as ‘hit’ trials and rewarded with a drop of water. Withholding licking in response to the no-go sound were counted as correct rejection trials (CR), and were not punished or reinforced. Licking in response to the no-go sound were false alarm trials (FA) that were followed by a mild punishment of white noise (duration of 3 seconds). Withholding licking for the go sound were counted as misses, and were not punished. The licking detector remained in a fixed and reachable position throughout the entire trial and mice were free to lick at any time. Licking before the response cue was allowed and did not lead to punishment or early reward. Note that the visual cue merely signals the start of the trial, but had no predictive power with respect to go or no-go condition.

### Training and performance

Nine mice were trained on the task. Mice were first handled and accustomed to head fixation before starting the schedule of water restriction. Before imaging began mice were conditioned to lick for reward after the go sound (presented within a similar trial structure as the task itself). Imaging began only after mice reliably licked for the presented sound (typically after the 1^st^ day; 200-400 trials). On the first day of imaging, mice were presented with the go sound for 50 consecutive trials, after which the no-go sound was gradually introduced (starting from 10% and increasing by 10% approximately every 50 trials). By the end of the 1^st^ day, the no-go sound reached 50% probability (Guo et al., 2014). During the 2^nd^ day, most mice continuously licked for both sounds. Thus, after roughly 100 trials, we increased no-go probability to 80% and waited for mice to perform three consecutive CR trials before returning to 50% probability. This was done for several times until mice increased their performance, specifically to learn to withhold licking for the no-go sound. In mice that still continued to lick for both sounds we also repeated the no-go sound several times until the mouse performed correctly. In all mice, a 50-50% protocol was reached typically on the 1^st^ or 2^nd^ day. Most mice learned the task within 3-7 days corresponding to roughly 1200-2000 trials (Fig. 1D). An effort was made to maintain a constant position of the mouse, speaker and cameras across imaging days in order to maintain similar stimulation and imaging conditions.

### Optical fiber setup

As mice trained on and performed the task we continuously imaged MGB population responses. A ferrule patch cable (M79L01; Thorlabs) was connected to the implanted cannula using a mating sleeve (ADAF1; Thorlabs) and the cable end was connected (SMA connector) to an optical imaging setup (FOM-2; MCI), enabling us to excite the MGB at 470 nm (blue; 0.3 mW power output kept constant throughout the experiments) and collect emission light at 520 nm (green). As a control, we also imaged mice during the task while exciting the MGB with a wavelength that does not excite the calcium indicator (565 nm).

#### Histology

Mice were given an overdose of Pental and were perfused transcardially with phosphate-buffered saline (PBS) followed by 4% paraformaldehyde (PFA) in PBS. Brains were post-fixed for 12–24 h in 4% PFA in PBS and then cryoprotected for >24 h in 30% sucrose in PBS. Then, 100 μm coronal slices of the entire brain were made using a freezing microtome (Leica SM 2000R), incubated for 15 min in 2.5 μg/ml of DAPI (4’,6-diamidino-2-phenylindole), mounted onto glass slides, and imaged using an Olympus IX-81 epi-fluorescent microscope with a 4× and 10× objective lens (0.16 and 0.3 NA; Olympus). Fiber tracks and GCaMP6f expression were detected in all mice and were mainly localized to the MGB (Fig. 1C and Supplementary Fig. 2).

#### Body tracking

In addition to fiber photometry, we tracked body movements of the mouse during the task (Fig. 1B and 5A). The mouse was illuminated with a 940-nm infra-red LED. A body camera monitored the movements of the mouse at 30 Hz (using a DMK 23UV024 camera). We used movements of both forelimbs and the head/neck region to assess body movements (Fig. 1A; see *Data Analysis* below). Mice performed the task in the dark.

#### Data analysis

Data analyses and statistics were performed using custom-written code in MATLAB. For MGB signals, each trial was normalized to baseline several frames before the visual cue (frame 0 division). We did not find a substantial difference in baseline fluorescence across imaging days and also within an imaging day, implying there was no over bleaching of the signal. This enabled us to quantitatively compare ΔF/F signals across long time periods. To better control for possible changes in MGB signal across days, we tested for frequency tuning (playing a range of frequencies 4-40 kHz; 10, 20, 30, 40 dB attenuations relative to 62 dB; duration 0.1 second) in each mouse, both before and after training. We did not see a significant difference in ΔF/F response between the cases (Fig. S5; P>0.05; Signed rank test).

Next, we divide trials based either on stimuli (i.e. go or no-go) or on choice (i.e. lick or no-lick). MGB ΔF/F signals were plotted in 2 dimensional temporal spaces where the x-axis is the trial temporal structure and the y-axis is the learning profile across trials and days (Fig. 2A). From this 2D temporal space we averaged across trials during the novice and expert phases (defined as the first and last 500 trials respectively; Fig. 2B). Alternatively, we averaged across time frames within the trial structure, to obtain an MGB response curve across learning (Fig. 4A; see below).

### Discrimination power between different trial types

To measure how well could neuronal populations discriminate between trial types (i.e. hit, miss, CR and FA), we calculated a receiver operating characteristics (ROC) curve between the distribution of a pair of trial types and calculated its area under the curve (AUC). We define three different AUC measures by comparing hit trials to either CR (task AUC; brown; i.e. different stimuli and different choices), to FA (stim AUC; magenta; i.e. different stimuli but similar choice) or to Miss (choice AUC; green; i.e. similar stimuli but different choice) trials (Fig. 3A). The AUC value ranges from 0 to 1 and quantifies the accuracy of an ideal observer. AUC values close to 0.5 indicate low discrimination whereas values near away from 0.5 indicate high discrimination. In addition, we calculated AUC values for each time frame along the trial for the novice and expert conditions separately (i.e. first and last 500 trials). To assess significance, we calculated the sample distribution by trial shuffling between go and no-go sounds (n=100 iterations). When the signal exceeded the mean±3std of the sample distribution it was defined as significant (Fig. 3B, arrows).

### Calculation of learning curve and MGB response curves

To calculate the learning curve for each mouse, trials were binned (n=50 trials with no overlap) across learning and the performance (defined as d’ = *Z*(hit/(hit+miss)) – *Z*(FA/(FA+CR)) where *Z* denotes the inverse of the cumulative distribution function) was calculated for each bin. Next, each behavioral learning curve was fitted with a sigmoid function

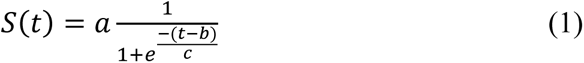

Where *a* denotes the amplitude, *b* the time point (in trial numbers) of the inflection point, and *c* the steepness of the sigmoid. A d’=1 was defined as the threshold and mice were ordered based on the trial number in which they crossed threshold (i.e. learning threshold; Fig. 1E). Different d’ thresholds did not change the order with which mice learned (see Fig. 1E).

To compare the behavioral learning curve with responses in the MGB we calculated the mean MGB response across learning (in the same 50 trial bins as the learning curve), averaged during late times (0.6-1 seconds after stimulus onset). Our main focus in this study was on the go sound (hit and miss trials grouped together). Therefore, stimulus identity was kept similar across learning. MGB response curves were also fitted with a sigmoid function (black curves in Fig. 4A; Curves were smoothed with a Gaussian kernel (2σ=9) for visualization only). The sigmoid fits of the MGB response curves were normalized between 0 and 1 in order to compare between curves from different mice. In addition, we calculated the MGB response curve for each time frame separately. This was correlated with the fixed learning curve of the mouse, to obtain a time course (within the trial) correlation coefficient value between the MGB response curve and the learning curve (Fig. 4B; no smoothing applied).

### Calculating body movements

We used a body camera to detect general movements of the mouse (30 Hz frame rate; Fig. 1A and 5A). For each imaging day, we first outlined the forelimbs and the neck areas (one area of interest for each), which were reliable areas to detect general movements. Next, we calculated the body movement (1 minus frame-to-frame correlation) within these areas as a function of time for each trial. Thresholding at 3 s.d. (across time frames before stimulus cue) above baseline resulted in a binary movement vector (either ‘moving’ or ‘quiet’) for each trial (Gilad et al., 2018; Gilad and Helmchen, 2019). This was done for each trial to achieve a 2D space of movement probability within the trial temporal structure (x-axis) versus the learning process (i.e. trial number; y-axis; Fig. 5B). To obtain MGB signals that are ‘movement-free’, i.e. do not contain direct effect of body movements, we detected the first movement onset for each trial, defined as 0.2 seconds before crossing the movement threshold (Fig. 5E). Next, MGB signals were truncated from movement onset and onwards in a single trial manner. This analysis resulted in MGB trials, each with a different length, that did not contain body movements (Fig. 5F,G). Similar results were obtained using other body parts such as the nose and mouth (including licking).

### Statistical analysis

In general, non-parametric two-tailed statistical tests were used. The Whitney rank sum test was used to compare between two medians from two populations and the Wilcoxon signed rank test was used to compare a population’s median to zero (or between two paired populations). Multiple group correction was used when comparing between more than two groups. Significance was set at p=0.05.

## Supplementary information

**Figure S1.**
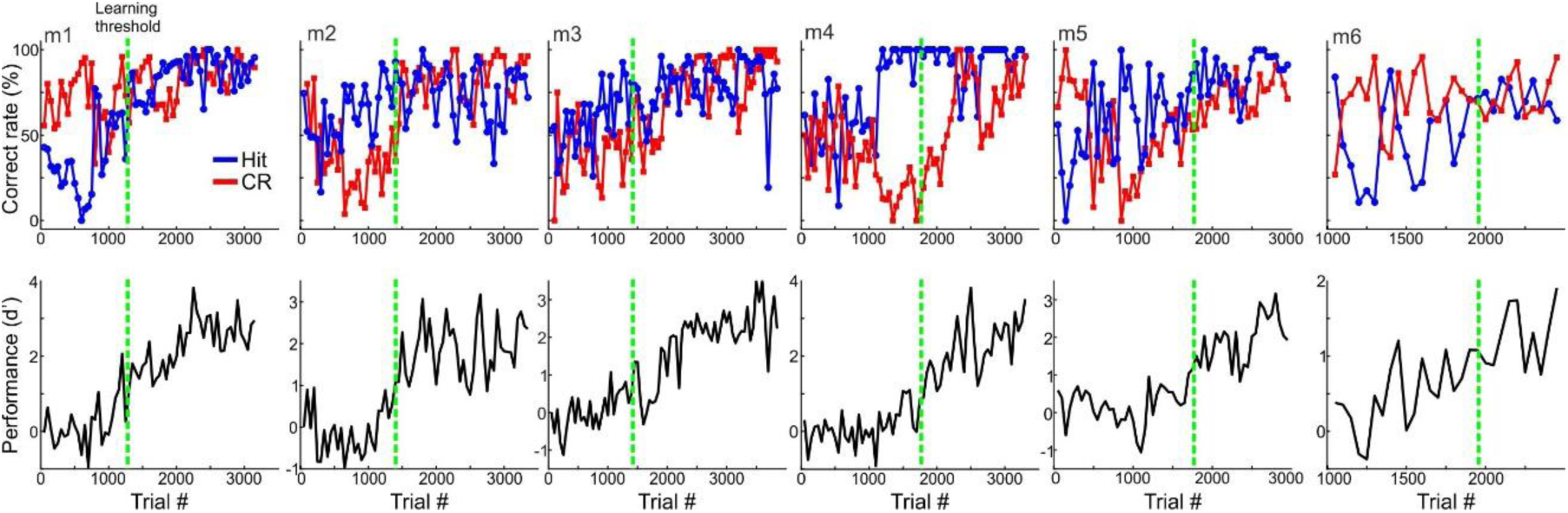
Behavioral performance and learning curves. Behavioral performance for all six mice, plotting the time course of Hit and CR rates in percent (top) and d’ (bottom) as a function of trial number. The learning threshold when mice reached d’ = 1 is indicated by vertical dashed green lines.

**Figure S2.**
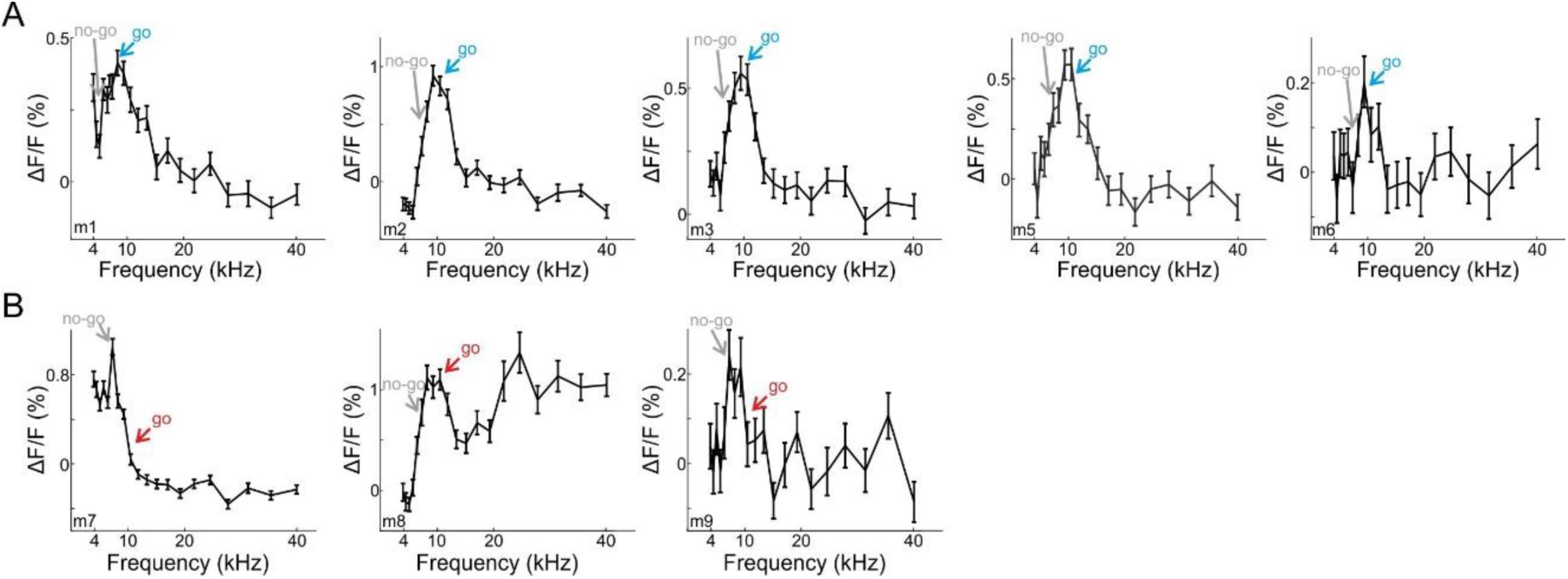
Frequency tuning for each mouse. **A.** Frequency tuning curve tuned for the go frequency. The frequencies used as go, and no-go are marked. Mouse 4 did not have a frequency curve. **B.** Frequency tuning curve tuned away from the go frequency.

**Figure S3.**
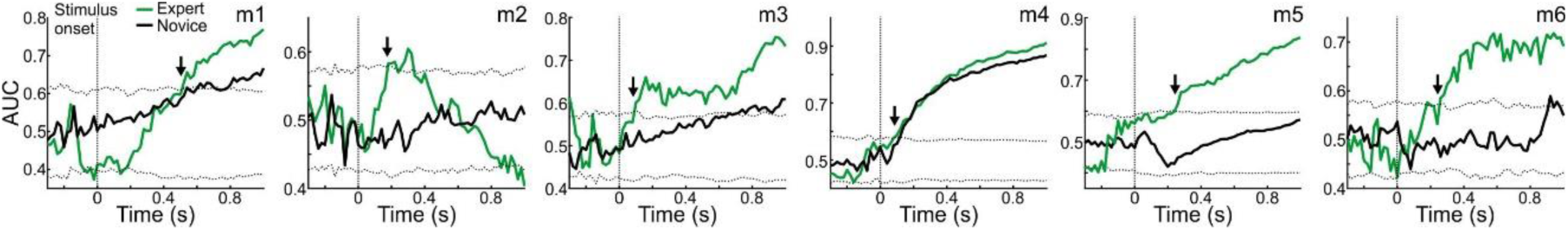
Choice-AUC responses for all mice. Choice AUCs during the trial for all mice during novice (black lines) and expert (colored lines). Similar to Figure 3B, right panel. Dashed black lines display the mean±3s.t.d of trials shuffled data. Arrows indicate the latency of the AUC measure (i.e. the first time frame exceeding the shuffled data).

**Figure S4.**
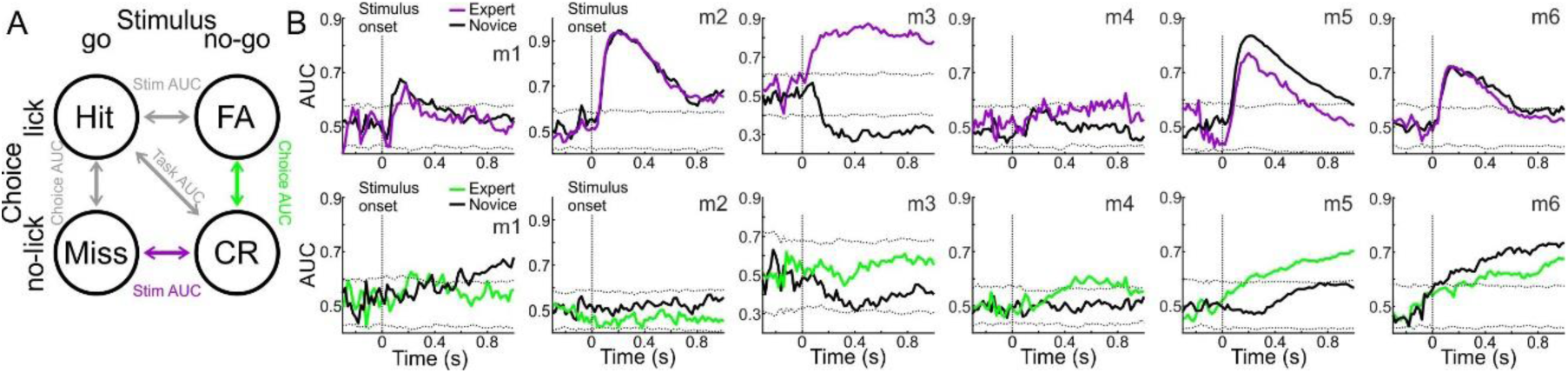
Stim-AUC and Choice-AUC based on other trial type pairs. **A.** Schematic of alternative single trial discrimination measures using different trial types: Miss vs. CR (purple; Stim AUC) or FA vs. CR (light green; Choice AUC). Stim-AUCs (top) and choice AUCs (bottom) during the trial for all mice during novice (black lines) and expert (colored lines; purple for stim-AUC; Light green for choice-AUC). Similar to Figure 3B. Dashed black lines display the mean±3s.t.d of trials shuffled data.

**Figure S5.**
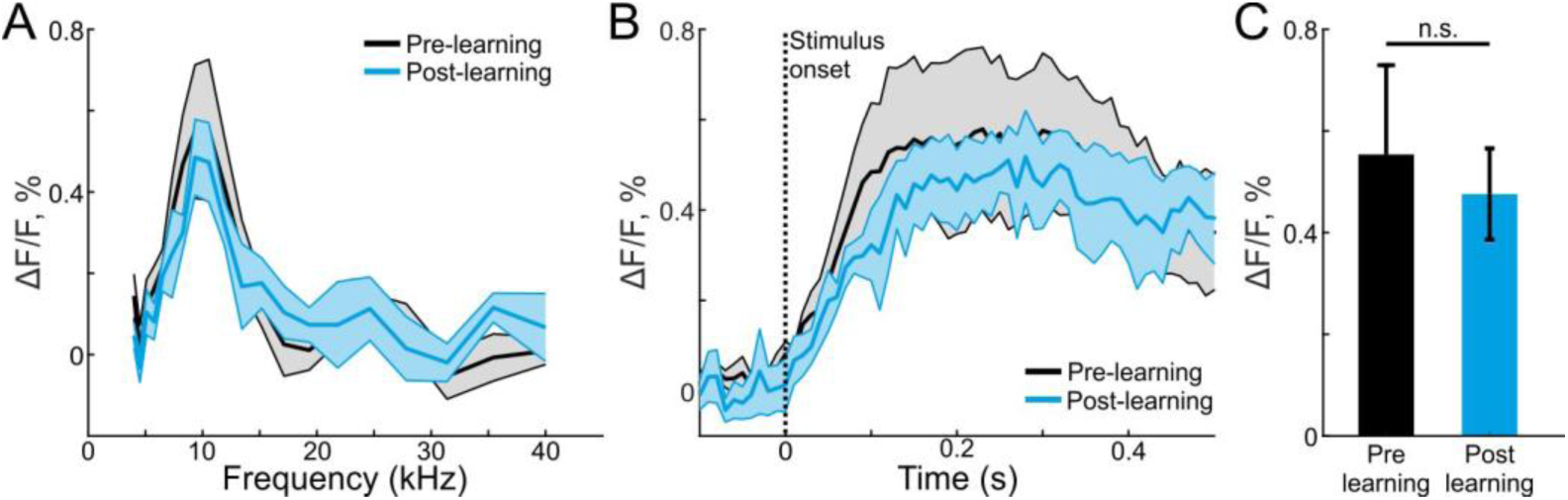
Tonotopic mapping is similar before and after learning. A. Frequency tuning curves (obtained during tonotopic mapping of passive listening) averaged across all mice before (blue trace) and after (black trace) learning. B. Average responses to the go frequency before (blue trace) and after (black trace) learning. C. Average evoked response (calculated from the 100-300 ms after stimulus onset) to the go frequency before (blue trace) and after (black trace) learning. Error bars are s.e.m across mice. n.s. – not significant. Wilcoxon sign rank test.

**Figure S6.**
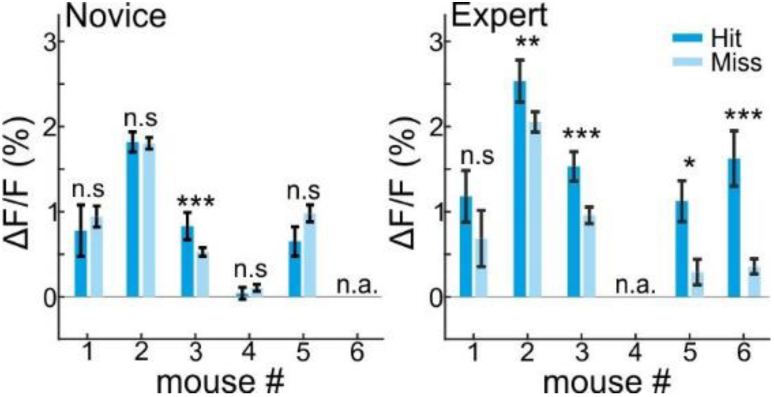
Movement-free MGB choice encoding in novice and expert mice. Mean MGB responses (averaged during stimulus presentation) for movement-free hit and miss trials per mouse during the novice (left) and expert (right) cases. Error bars are s.e.m across trials. *P < 0.05. **P < 0.01. ***P < 0.001. n.s. – not significant. Wilcoxon rank sum test.

**Figure S7.**
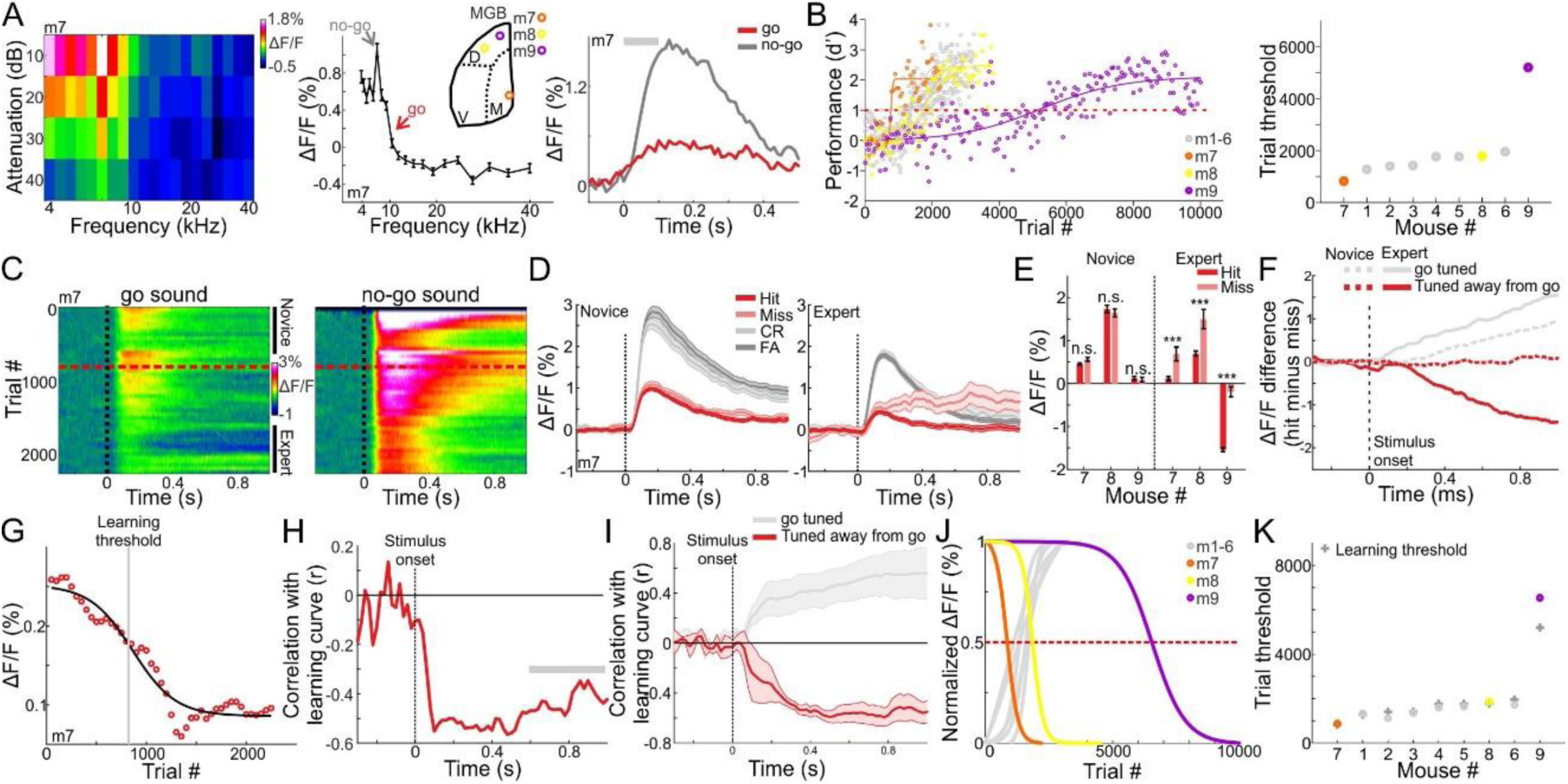
MGB responses tuned away from the go frequency are suppressed during learning. **A.** *Left:* An FRA plot (attenuation versus frequency) from one example recording of mouse #7. *Middle:* Frequency tuning curve for the same example mouse with peak tuning frequency away from the go sound. *Right:* Average responses to the go (red) and no-go (gray) sounds. **B.** *Left:* Behavioral learning curves for all mice (n=3) tuned away from the go frequency depicting the performance (d’) as a function of trial number. Each learning curve was fitted with a sigmoid function. Dashed red line indicates the performance threshold (d’=1). In gray are the go preferring mice similar to Fig. 1D. *Right:* Learning threshold (the trial number where the learning curve crossed the performance threshold) for all mice. In gray are the go preferring mice similar to Fig. 1E. **C.** 2-dimensional plots of the calcium responses in MGB during the trial (x-axis) and across learning (y-axis; 50 trial bins) for one example mouse divided into go (left) and no-go (right) sounds. Dashed black line indicates stimulus onset (1 s duration) and dashed red line indicates the learning threshold. **D.** Calcium responses when the mouse was novice (left) and expert (right). Traces are shown separately for different trial types (hit, miss, FA and CR trials; same data as in ‘C’). Shaded error bars are s.e.m across trials. **E.** Mean calcium response during stimulus presentation per mouse in hit and miss trials. Left-novice mice, Right-expert mice. Error bars are s.e.m across trials. ***P < 0.001. n.s. – not significant. Wilcoxon rank sum test. **F.** Choice responses, averaged across all three mice for the expert (solid line) and novice (dashed line) conditions. Gray traces are for go preferring mice, same as in Fig. 2D. **G.** MGB response curve along learning in one mouse (compare to Fig. 4A). H. Correlation between MGB response curves and learning curves as a function of time for an example mouse that is tuned away from the go frequency. **I.** Correlation between MGB response curves and learning curves as a function of time, averaged across the 3 mice. Error bars are s.e.m across mice. Gray traces are for go preferring mice, same as Fig. 4C. **J.** Normalized MGB curve fits of mice tuned away from the go frequency. Dashed red line indicates threshold. Gray traces are for go preferring mice, same as Fig. 4D. **K.** Maximal change in MBG responses (i.e. the trial number in which the MGB fit crossed 0.5, dots) as a function of learning threshold (gray crosses). Gray circles are for go preferring mice, same as Fig. 4E.

